# GABA, glucose and insulin orchestrate CD4^+^ T cells effector functions

**DOI:** 10.1101/2023.04.18.537327

**Authors:** Z Jin, H Hammoud, AK Bhandage, SV Korol, O Trujeque-Ramos, S. Koreli, AI Chowdhury, FA Sandbaumhüter, ET Jansson, PE Andrén, P Bergsten, M Kamali-Moghaddam, B Birnir

## Abstract

Metabolic programs of immune cells are closely linked to their effector functions. Physiological molecules including glucose, insulin and γ-aminobutyric acid (GABA) provide environmental cues and guidance, although how they coordinate to regulate the cells is still being unraveled. Here, we demonstrate that GABA-mediated reduction of metabolic activity and release of inflammatory molecules, including IFNγ and IL-10, was abolished in human CD4^+^ T cells when the glucose concentration was elevated above normal levels. Insulin enhanced the GABA_A_ receptors-mediated currents and Ca^2+^ influx. GABA decreased, whereas insulin sustained glycolysis but in a SGLT (Na^+^-glucose transporter)-dependent manner. In high glucose (16.7 mM), the SGLTs antagonist phlorizin alone or together with GABA restored the inhibition of IFNγ and IL-10 release. This study exposes concerted effects of GABA, glucose and insulin on CD4^+^ T cells metabolic activity and release of inflammatory molecules, and identifies a role for SGLTs in CD4^+^ T cells function.

## Introduction

Human T cells are a crucial component of the adaptive immune response and provide essential immune protection throughout life. Upon antigen stimulation, naïve T cells undergo clonal expansion and further differentiate into effector cells. During this process, metabolic reprogramming of T cells occurs and is intricately linked to their effector functions^1^. A variety of soluble molecules including amino acids, other nutrients and hormones that are present in blood, lymph nodes and tissue microenvironment influence T cell function^2–4^. γ-aminobutyric acid (GABA), is primarily known as an inhibitory neurotransmitter in the central nervous system, but is increasingly recognized as an immunomodulatory molecule^5–7^. We and others have previously shown that GABA inhibits proliferation and cytokine secretion by activating GABA_A_ receptors in CD4^+^ T cells^8–10^. The GABA concentration in the plasma from healthy subjects is around 500 nM and increases somewhat in patients with Type 1 diabetes (T1D) or major depression^10,11^. Glucose is the primary source of energy for activated T cells but is also utilized for synthesizing biomolecules to support rapid proliferation and differentiation of the cells^12,13^. At the time of activation, glucose is transported by facilitated glucose transporters that equilibrate glucose across the cell membrane^3^. The blood glucose concentration varies under physiological (around 5.6 mM) and pathophysiological conditions, for example, in diabetes-related hypoglycemia (< 3 mM) or hyperglycemia (> 11 mM)^14^. In human clinical trials, GABA treatment has been shown to be beneficial for glycemic control in T1D individuals^15,16^. Insulin, a hormone mainly associated with glucose homeostasis^14^, also influences T cell function by modulating their metabolism^17^ We and others have previously demonstrated that GABA signaling regulates hormone secretion in human islets^18,19^. However, how GABA, glucose and insulin orchestrate effective immune activities of T cells is still being uncovered.

In the present study, we observe that, in human CD4^+^ T cells, increasing the glucose concentration above normal levels attenuates GABA-mediated inhibition of metabolic activity and release of inflammatory proteins. Furthermore, insulin augments GABA_A_ receptors-mediated currents and Ca^2+^ influx in a manner that is dependent on the glucose concentration. We also reveal the presence of functional Na^+^-glucose transporter 2 (SGLT2) in activated CD4^+^ T cells and, thereby, identify the possibility that T cells can raise the intracellular above the environmental glucose concentration. Finally, we show that GABA, insulin, glucose and an SGLT2 inhibitor differentially modulate glycolysis and release of IFNγ and IL-10 in activated CD4^+^ T cells. These results demonstrate coordinated influence of GABA, glucose and insulin on metabolic activity and release of inflammatory proteins, and highlight the involvement of SGLTs in CD4^+^ T cell functions.

## Results

### GABA inhibition of activated T cell function is affected by glucose

Physiological blood glucose concentration 2h after a meal, is normally maintained around 5.6 mM in healthy individuals, while it is higher in diseases such as diabetes (> 7 mM)^14^ or Covid-19^20^. Metabolic activity of T cells is closely related to their function^1^. The physiological GABA concentration in human plasma is about 500 nM^10,21^ and it provides a homeostatic inhibition on T cells^10^. The culture media (e.g., RPMI1640) commonly used for *in vitro* T cell assays usually contain high glucose concentration (> 10 mM). Therefore, we examined effects of four glucose concentrations on the GABA inhibition of metabolic activity of CD4^+^ T cells (N=35–53 human donors; Fig. 1a; Supplementary Table 1, Supplementary Fig. 1). At glucose concentrations higher than the physiological level (10 mM and 16.7 mM), the inhibition by GABA (500 nM) was abolished (Fig. 1a). In absence of GABA, the metabolic activity of activated CD4^+^ T cells varied somewhat for the different glucose concentrations (5 mM vs. 2.8 or 10 mM; Supplementary Fig. 2a). When the GABA responses were compiled based on low-normal (2.8 mM, 5.6 mM) or high (10 mM, 16.7 mM) glucose concentrations, a clear shift was observed for GABA’s inhibitory response (Fig. 1b). At the level 0.9 or lower metabolic activity (Fig. 1b), cells from 74% of the donors were inhibited by GABA in the low-normal glucose group. The metabolic activity of cells from these same donors were not inhibited at the higher glucose concentrations (Fig. 1c). In contrast, cells from the remaining 26% donors were not inhibited by GABA at any glucose concentration (Supplementary Fig. 2b), in agreement with previous report ^10^. Intrigued by these results, we sought whether glucose affected other immunological functions regulated by GABA. Since glycolysis feeds substrates into anabolic metabolism, we studied if glucose modulated the GABA-inhibition of release of biomolecules from CD4^+^ T cells from the same donors (N=17) cultured in low-normal or high glucose. In each sample, 92 inflammation-related biomolecules commonly associated with inflammation were measured using Olink Target Inflammation protein panel, which is based on the multiplex proximity extension assay (PEA) technology. A total of 59 different proteins were detected. Similar to the results from the metabolic activity, the GABA-inhibition of proteins released from CD4^+^ T cells was glucose concentration-dependent (Fig. 1d). The GABA inhibition was optimal at the physiological glucose concentration (5.6 mM), where GABA decreased the release of 37 proteins. In contrast, GABA did not affect release of proteins in neither 10 mM nor 16.7 mM glucose (Fig. 1d). ELISA confirmed the PEA results for two cytokines; interferon gamma (IFNγ), the principal cytokine of CD4^+^ T helper1 cells and interleukin-10 (IL-10), the principal cytokine for CD4^+^ T helper2 and T regulatory cells (Fig. 1e, f). Further, in absence of GABA, increasing the glucose concentration alone enhanced release of IFNγ (Supplementary Fig. 2c) but not IL-10 (Supplementary Fig. 2d). Thus, GABA-dependent regulation of CD4^+^ T cells is linked to the extracellular glucose concentration.

**Figure 1.**
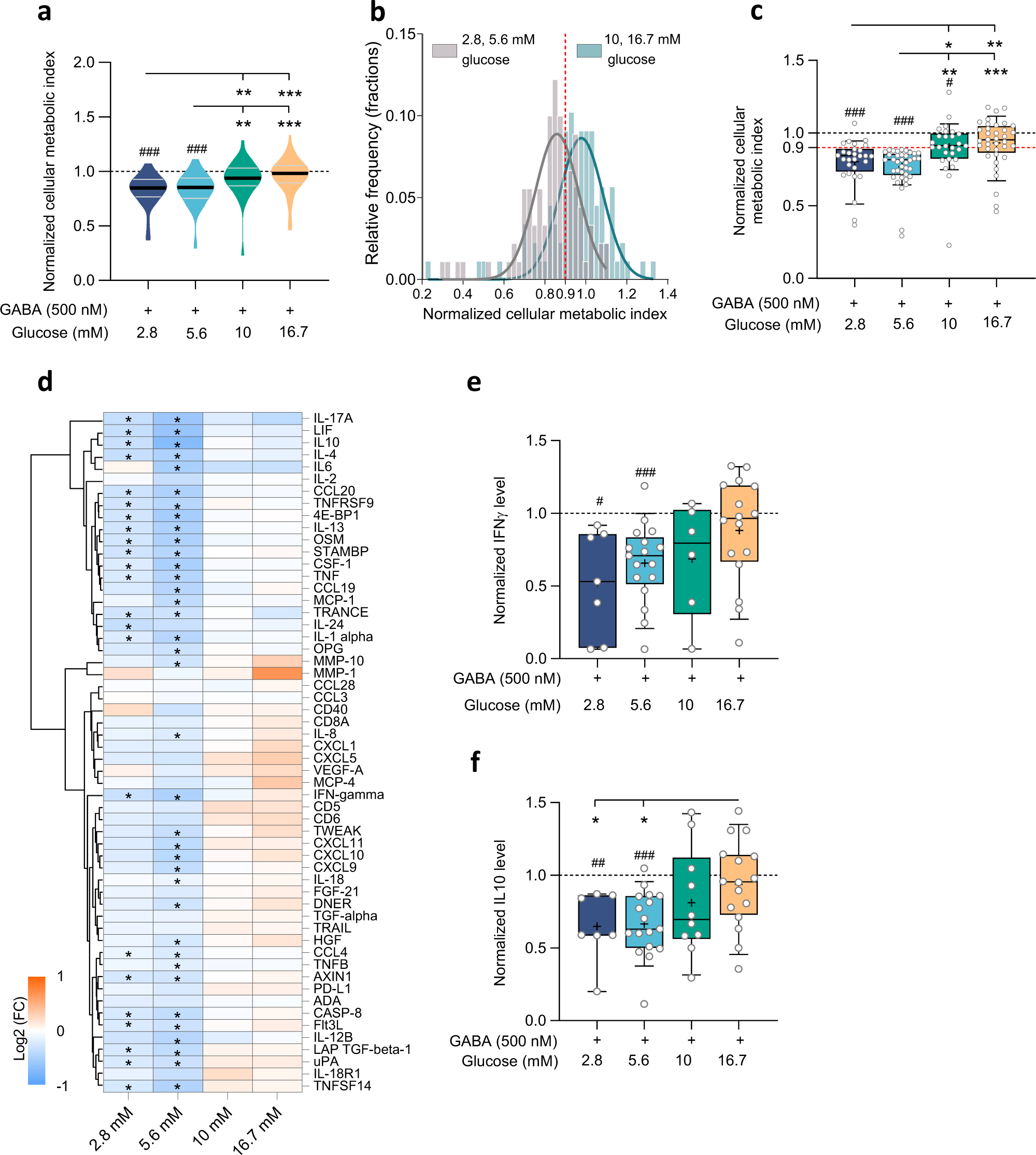
GABA effects on metabolic activity and cytokine release are glucose concentration-dependent in activated CD4^+^ T cells. **a**, Violin plots show cellular metabolic activity of activated CD4^+^ T cells as measured by MTT assay, in the presence of GABA at different glucose concentrations: 2.8 (N=36), 5.6 (N=54), 10 (N=35) and 16.7 mM (N=53). Horizontal black and grey lines indicate the median and quartiles, respectively. **b**, Distribution (histogram and Gaussian fit) of normalized cellular metabolic activity of CD4^+^ T cells at low-normal (2.8 and 5.6 mM) and high (10 and 16.7 mM) glucose concentrations. Solid lines: Gaussian fits, vertical broken line: 10% inhibition by GABA (500 nM). X-axis: normalized cellular metabolic index, Y-axis: relative frequency represented as fractions. **c**, Box plots show metabolic activity of activated CD4^+^ T cells from donors where ≥10% GABA inhibition was observed at 5.6 mM glucose; 2.8 (N=28), 5.6 (N=37), 10 (N=27) and 16.7 (N=36). **d,** Heatmap and hierarchical clustering of inflammatory-related biomolecules by PEA 72 h post-stimulation of CD4^+^ T cells, at different glucose concentrations (N=17). Data represent the mean log2 (fold change) for samples cultured in presence and compared to absence of GABA. **e, f,** Box plots show GABA inhibition of IFNγ **(e)** and IL-10 **(f)** release, as measured by ELISA, glucose concentrations (mM): 2.8 (N=7;7), 5.6 (N=17;17), 10 (N=6;10) and 16.7 (N=16;16) in **e** and **f**, respectively. Data were normalized to values of activated cells in the absence of drugs at each glucose concentration (**a, c, e, f**). Box and whisker (**c, e, f**): 10-90 percentiles, means: +, medians: black lines in the boxes. Statistics: one sample t-test when compared to activated cell group (#, p < 0.05; ##, p < 0.01; ###, p < 0.001), paired groups by repeated measures ANOVA or fitting a mixed-effects model, or ordinary one-way ANOVA followed by Tukey (**a, c, e, f**) for multiple comparisons (*, p < 0.05; **, p < 0.01; ***, p < 0.001). Statistical significance in **d (**p ≤ 0.05 and adjusted p-value q < 0.12) are marked with *. **N**: donors.

### Insulin enhances the GABA-activated currents and modulates GABA_A_ receptor expression in CD4^+^ T cells

Glucose homeostasis is influenced by a variety of hormones, with insulin being one of the key players. Interestingly, at least in excitable cells, insulin is also a regulator of GABA signaling^22–24^. The insulin receptor is not expressed in resting T cells but is prominent at 48 h post-activation^17^. Therefore, after 48-h activation, we incubated CD4^+^ T cells with insulin at physiologically relevant concentration (3 nM) and where unspecific activation of other receptors-types does not take place^17,24^. GABA activates two types of receptors in cell membranes; the GABA_A_ receptor (GABA_A_R), a pentameric chloride ion channel, that opens when GABA binds to the receptor and the GABA_B_ receptor (GABA_B_R), a G-protein coupled receptor, that is formed as a dimer. To-date, only one of the two GABA_B_ isoforms required for function has been detected in immune cells, whereas several GABA_A_ receptor subunits are identified in T cells^4,10^. We measured the GABA-activated GABA_A_R currents in intact activated CD4^+^ cells using perforated patch-clamp electrophysiology (Fig. 2a, b, c, d). Since the intracellular environment is unchanged, intracellular signaling that may take place is expected to function normally. The 500 nM GABA-activated GABA_A_R-mediated currents were blocked by the GABA_A_R antagonist picrotoxin. Picrotoxin is a specific, open channel blocker of GABA_A_ receptors^25^. These maximal current responses were only enhanced by insulin at 5.6 mM but not at 16.7 mM glucose (Fig 2a), but the apparent GABA affinity (EC_50_) for GABA_A_R was similar in 5.6 mM and 16.7 mM glucose, 0.7 nM and 0.9 nM GABA, respectively (Fig. 2b). Insulin then shifted the EC_50_ values more than 10-fold, to 0.04 nM and 0.06 nM GABA (Fig. 2c), for the two glucose concentrations, respectively. These results suggest that insulin increases the GABA_A_R plasma membrane surface number in 5.6 mM glucose but enhances the GABA affinity of GABA_A_R in CD4^+^ T cells at both glucose concentrations. The GABA_A_R agonists muscimol and TACA evoked currents similar to GABA (Fig. 2d). The currents reversed at −37 mV (Supplementary Fig. 3a), close to the calculated reversal potential for chloride (E_Cl_ = −39 mV). By regulating the membrane potential these very sensitive GABA_A_R channels may affect activity of other ion channels or flow of ions across the cell membrane^26^.

**Figure 2.**
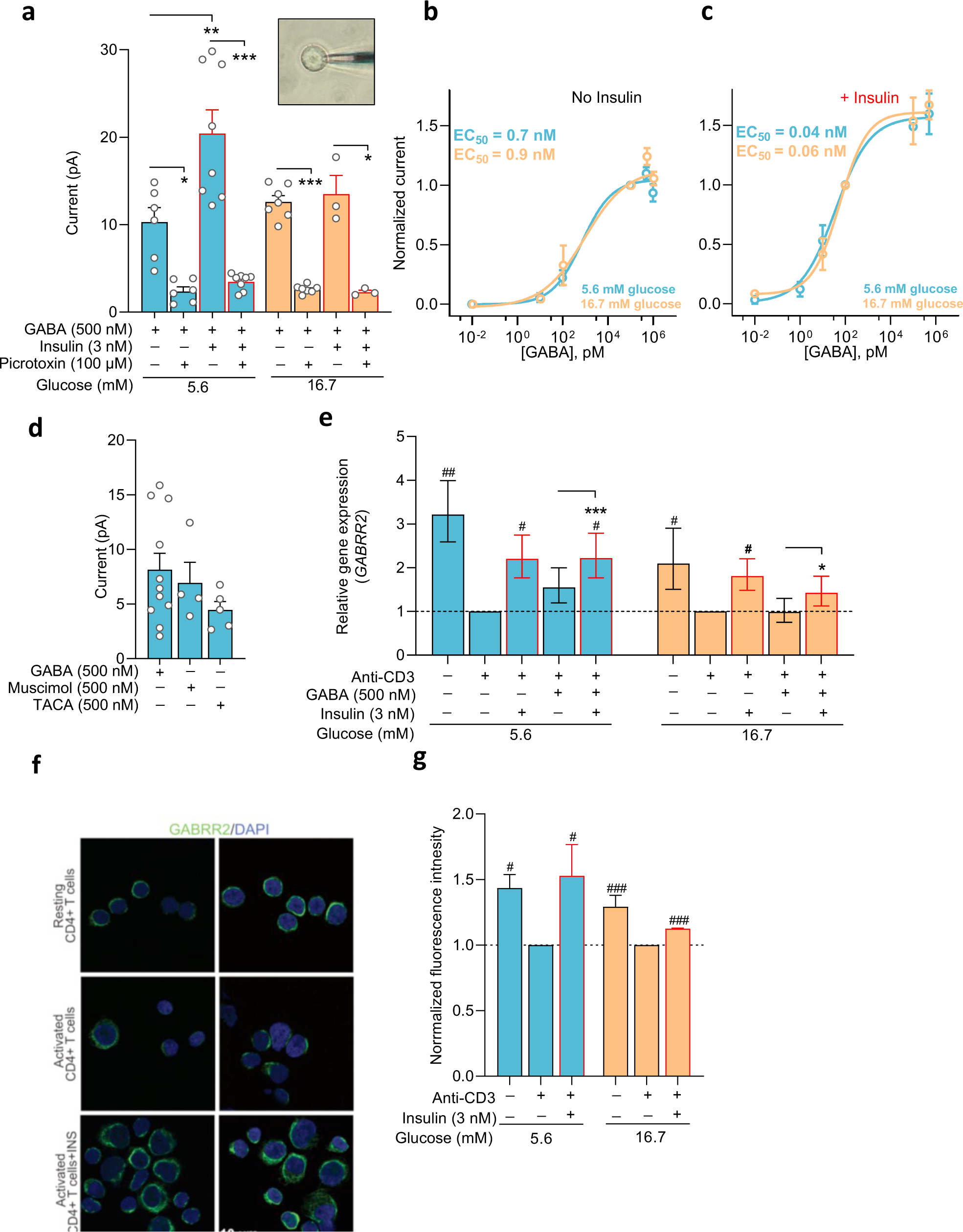
Insulin enhances GABA-activated GABA_A_R currents and regulates GABA_A_ receptor expression in activated CD4^+^ T cells. **a,** GABA-activated whole-cell currents (at +30 mV) in the absence or presence of a GABA_A_R antagonist picrotoxin (PTX) in activated CD4^+^ T cells without or treated with insulin (n=3-8, N=11). Insert: a patch-pipette attached to an activated CD4^+^ T cell. **b, c,** GABA dose-response curves and the calculated half maximal effective concentration (EC_50_) in cells without (**b**) or treated with (**c**) insulin. The currents were normalized to 100 nM (**b**) or 100 pM (**c**) GABA responses (n=3–11, N=27). **d,** Whole-cell currents (at +30 mV) evoked by GABA_A_R agonists GABA (n=11, N=9), muscimol (n=4, N=3) or TACA (n=5, N=4), in activated CD4^+^ T cells in 5.6 mM glucose. **e,** Relative expression of *GABRR2* mRNA in resting and anti-CD3 antibody-activated cells in 5.6 or 16.7 mM glucose (N=6) analyzed by 2^-ΔΔCt^ method. **f,** Representative images from immunofluorescence co-staining of GABRR2 (green) and DAPI (blue) in CD4^+^ T cells in 5.6 or 16.7 mM glucose. **g,** Bar graphs show quantification of GABRR2 fluorescence intensity in CD4^+^ T cells (n>50 cells/group, N=3). Data represent mean ± SEM in (**a, d, g**) and geometric mean ± SEM in (**e**). Data were normalized to values of activated cells in the absence of drugs at each glucose concentration (**e, g**). Curve fitting (**b, c**): nonlinear regression with sigmoidal polynomial (4PL) model. Statistics: ordinary one-way ANOVA followed by Tukey (**a, d, g**) or repeated measures ANOVA followed by Fisher’s LSD test (**e**) for multiple comparisons. (*, p < 0.05; **, p < 0.01; ***, p < 0.001. #, p < 0.05; ###, p < 0.001when compared to activated cell group). **n**: cells, **N**: donors.

Among the 19 GABA_A_R subunits, the GABA_A_R ρ2 (*GABRR2*) is a highly expressed subunit in peripheral blood mononuclear cells^10^. We therefore examined the ρ2 mRNA and protein expression levels in CD4^+^ T cells. Irrespective of glucose concentration, activation of CD4^+^ T cells decreased the expression of the *GABRR2* transcript relative to resting cells, whereas insulin increased the expression of the *GABRR2* (Fig. 2e). In CD4^+^ T cells fluorescent labelling of the GABRR2 was detected by specific antibody (Fig. 2f, g). Taken together, the data show that insulin potentiates GABA signaling in activated CD4^+^ T cells by enhancing ρ2-containing GABA_A_R expressed in the cells.

### Insulin enhances GABA-activated Ca^2+^ entry in CD4^+^ T cell

The cytoplasmic concentration of Ca^2+^ ions is tightly regulated in T cells but can be increased by release from intracellular stores or by entry through plasma membrane Ca^2+^ ion channels^27^. A representative live cell-Ca^2+^ recording trace measured from activated CD4^+^ T cells (Fig. 3a) showed that acute GABA (500 nM) application induced transient calcium signal, which was, in fact, influx of extracellular calcium and not due to release from intracellular stores as no signal could be detected in absence of extracellular calcium (Fig. 3a). Analyzed live cell-Ca^2+^ recording data from four donors is shown in Fig. 3b. Transient change in glucose from 5.6 to 16.7 mM did not alter the 500 nM GABA-activated Ca^2+^ response (Supplementary Fig. 3b). Insulin incubation of cells enhanced the physiological saturating (500 nM) GABA-induced Ca^2+^ signals in 5.5 mM but not in 16.7 mM glucose (Fig. 3c), whereas no differences were observed for baseline Ca^2+^ levels in cells with or without insulin incubation (Supplementary Fig. 3c). The GABA-activated Ca^2+^ signal was blocked by the GABA_A_R antagonist picrotoxin (Fig. 3d), while no effect was observed by the GABA_B_R antagonist, CGP52432 (Supplementary Fig. 3d). This prompted us to examine dose-response relationship between GABA and the Ca^2+^ signals in cells with or without insulin incubation (Fig. 3e, f, g; Supplementary Fig. 3e). Representative time-lapse recording of a CD4^+^ T cell, 16.7 mM glucose, with insulin, demonstrates that raising GABA concentrations induced increasing Ca^2+^ signals (Fig. 3e). Insulin incubation of the cells lowered the EC_50_ values of the GABA-activated Ca^2+^ signals. The apparent GABA affinity of the Ca^2+^ signal was 40 nM and 133 nM GABA in 5.5 mM and 16.7 mM glucose, and shifted more than 1000-fold to 0.01 nM and 0.086 nM GABA with insulin, respectively (Fig. 3f, g). The GABA-induced Ca^2+^ response and enhancement by insulin is in accordance with our electrophysiological patch-clamp data (Fig. 2a, b, c). These data demonstrate that physiological GABA (pM – nM) increases Ca^2+^ entry, and the response is profoundly enhanced after insulin treatment.

**Figure 3.**
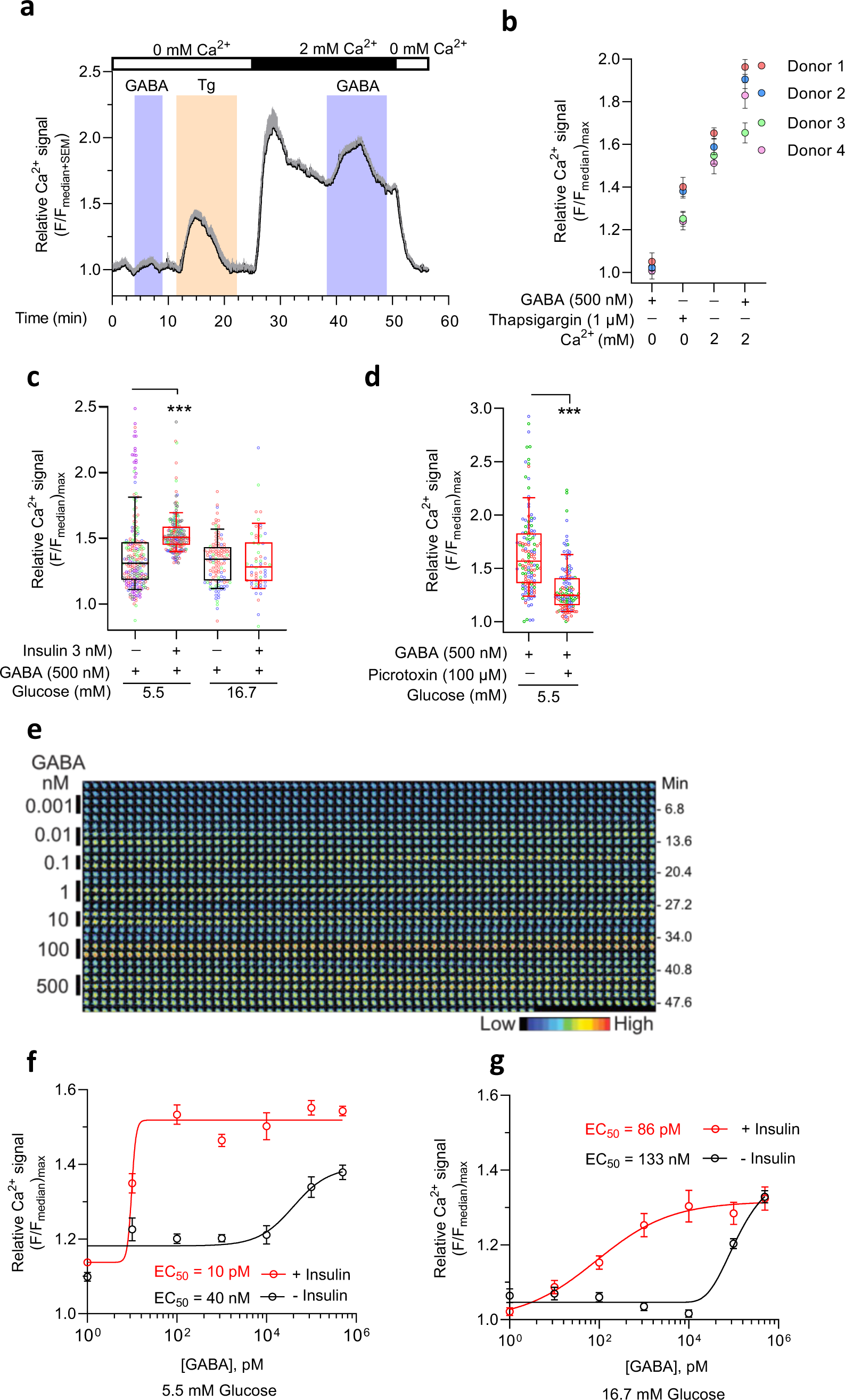
Insulin elevates GABA-activated Ca^2+^ entry in human CD4^+^ T cells. **a,** Mean Ca^2+^ signal intensity trace (black line) and SEM (gray whiskers) recorded 72 h after activation of cells (n=14). Colored bars: Perfusion time of drugs (GABA 500 nM and thapsigargin, Tg 1 μM) in 0 or 2 mM Ca^2+^-containing perfusion medium. **b**, Relative Ca^2+^ signal (mean ± SD) recorded in activated CD4^+^ T cells (n=14, 20, 41, 47, N= 4). **c,** GABA-activated relative Ca^2+^ signal recorded in activated CD4^+^ T cells (n=58-232, N=3-5) without or treated with insulin. **d,** GABA-activated relative Ca^2+^ signal in activated CD4^+^ T cells (n=131, N=3) in absence or presence of picrotoxin, with insulin. **e,** Representative time-lapsed micrographs of a live Ca^2+^ imaged cell (16.7 mM glucose, with insulin). GABA concentrations (vertical bars on the left) were sequentially perfused (time indicated as min on the right) and inter-spaced with medium applications. Color scale: Relative Ca^2+^ fluorescence intensity. **f, g,** Dose-response relationship and the calculated half maximal effective GABA concentration (EC_50_) for GABA-induced Ca^2+^ signals in activated CD4^+^ T cells without or treated with insulin in 5.5 mM (**f**) or 16.7 mM (**g**) glucose (n=63-279, N=3-8), see also Supplementary Figure 3e. In (**c, d**), individual data point represents relative Ca^2+^ signal recorded from each cell and is color-coded based on donors and box-whisker plots display the 10-90 percentile at the whiskers. Statistics: Two-tailed unpaired (**c**) or paired (**d**) Student’s t-test, ***, p < 0.001. **n**: cells, **N**: donors.

### GABA_A_R mediates GABA-activated Ca^2+^ entry in CD4^+^ T cells

To confirm the GABA_A_R connection in the evoked Ca^2+^ signal, we further examined the Ca^2+^ response to GABA_A_R agonists and antagonists. The application of GABA_A_R agonists, muscimol (Fig. 4a) or TACA (Fig. 4b) increased relative Ca^2+^ signals, which were inhibited by GABA_A_R antagonists, TPMPA (Fig. 4c) or picrotoxin (Fig. 4a, b, c), in activated CD4^+^ T cells treated with or without insulin. TACA and TPMPA are more selective for π-containing GABA_A_R^25,28,29^. Ca^2+^ release-activated Ca^2+^ (CRAC) channels mediate store operated calcium entry (SOCE) in T cells that is central to the Ca^2+^ homeostasis and cytoplasmic Ca^2+^ levels^27^. The GABA-activated Ca^2+^ signal was inhibited by 1 μM YM48583 (Fig. 4d), an antagonist of SOCE, revealing the Ca^2+^ ion channel participating in the cascade. The tight scaling of the Ca^2+^ signal from GABA_A_R activation to Ca^2+^ entry and the sensitization by insulin identifies a specific, focused mechanism regulating Ca^2+^ signaling in CD4^+^ T cells.

**Figure 4.**
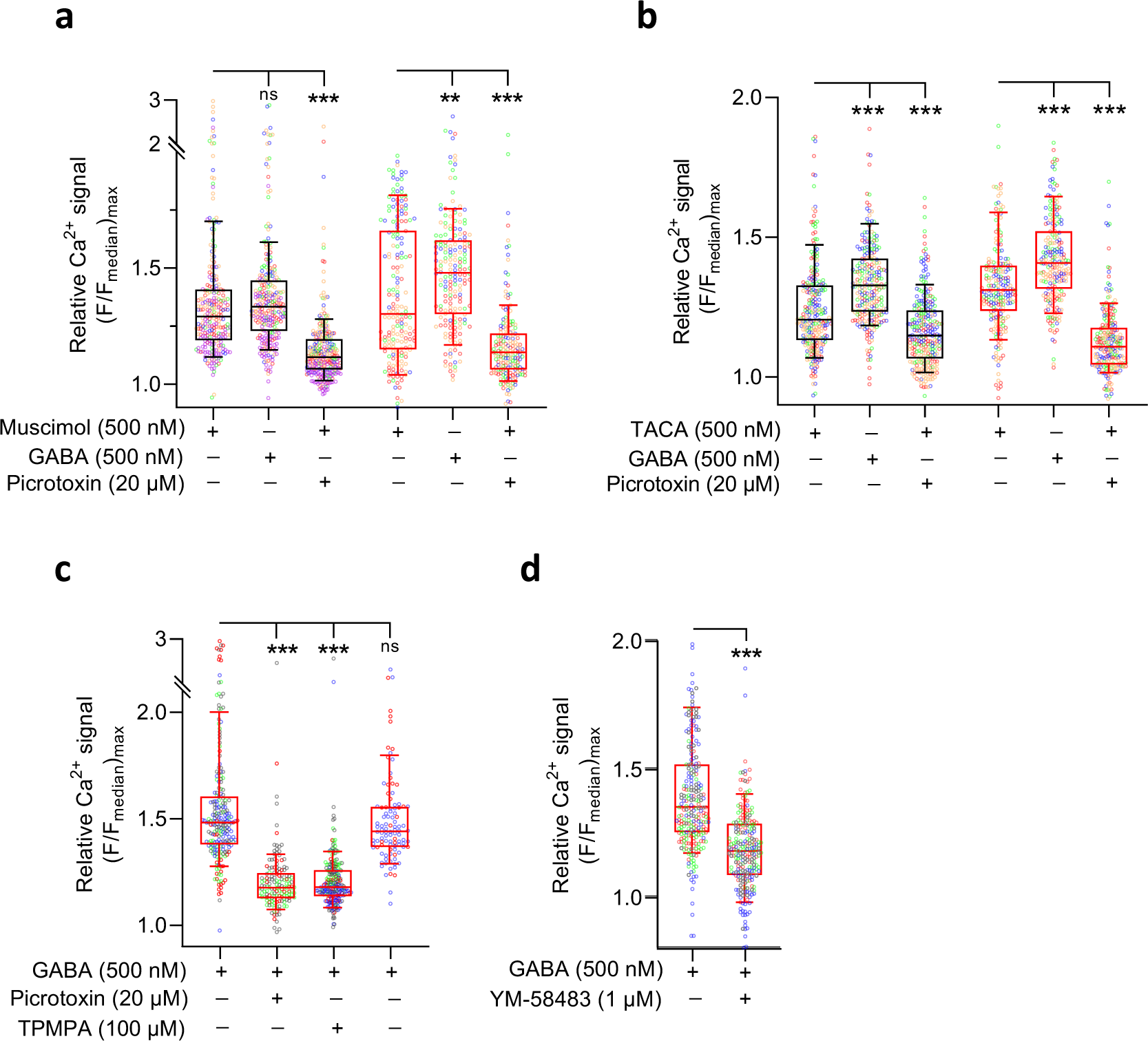
Pharmacological modulation of GABA-activated Ca^2+^ entry in human CD4^+^ T cells. Effects of GABA_A_R agonist, muscimol (500 nM, **a**) and TACA (500 nM, **b**), GABA_A_R antagonist picrotoxin (PTX, 20 μM, **a, b, c**) and TPMPA (100 μM, **c**), and SOCE antagonist YM48583 (1 μM, **d**) on relative Ca^2+^ signal in activated CD4^+^ T cells in 5.5 mM glucose (n=191-265, N=3-5). Individual data point represents relative Ca^2+^ signal recorded from each cell and is color-coded based on donors. Box-whisker plots (black without and red with insulin) display the 10-90 percentile at the whiskers. Statistics: repeated measures ANOVA followed by Tukey multiple comparison test (**a, b, c**) or two-tailed paired (**d**) Student’s t-test. ns, p > 0.05, **, p < 0.01, ***, p < 0.001. **n**: cells, **N**: donors.

Insulin is known to regulate intracellular signaling and gene expression, whereas less is known about effects of GABA. However, out of the nine examined genes encoding transcription factors or kinases, only *NFATC2* (nuclear factor of activated T cells 2) was significantly reduced by GABA at 5.6 mM glucose (Supplementary Table 2). NFATs proteins are activated by increased Ca^2+^ in the cytoplasm and translocate to the nucleus, where they regulate transcription of many proteins including inflammatory molecules^30^.

### GABA regulation of glycolysis is glucose-dependent

That GABA regulated expression of signaling pathways and proteins was revealed by mass spectrometry (MS) of activated CD4^+^ T cells cultured in the presence or absence of GABA (Fig. 5a, Supplementary Table 3). Hexokinase 1 (HK1), the first enzyme in the glycolytic pathway, was significantly down-regulated by GABA in both 5.6 mM and 16.7 mM glucose (Fig. 5b). HK activity was negligible under resting conditions but increased about 60 times in activated CD4^+^ T cells in both 5.6 mM and 16.7 mM glucose. Intriguingly, GABA decreased the HK activity only at 5.6 mM glucose and insulin had no further effect (Fig. 5c). T cells switch to aerobic glycolysis upon activation as their predominant way to generate adenosine 5’-triphosphate (ATP) and to accumulate biomass^31,32^, the so-called Warburg effect. It results in glucose-derived pyruvate being converted to lactate that can be detected by measuring extracellular acidification rate (ECAR)^33^. Since HK is the gate-keeper for the glycolytic pathway, we examined if glucose, GABA or insulin modulated glycolysis. Representative ECAR traces and average values for activated CD4^+^ T cells at 5.6 mM and 16.7 mM glucose in the presence or absence of GABA and insulin are shown in Fig. 5d and e. GABA but not insulin inhibited glycolysis and glycolytic capacity but only in 5.6 mM glucose. No effect of GABA at higher glucose suggested the possibility that either the intracellular glucose might affect the GABA inhibition of glycolysis or there were other competing mechanisms at work.

**Figure 5.**
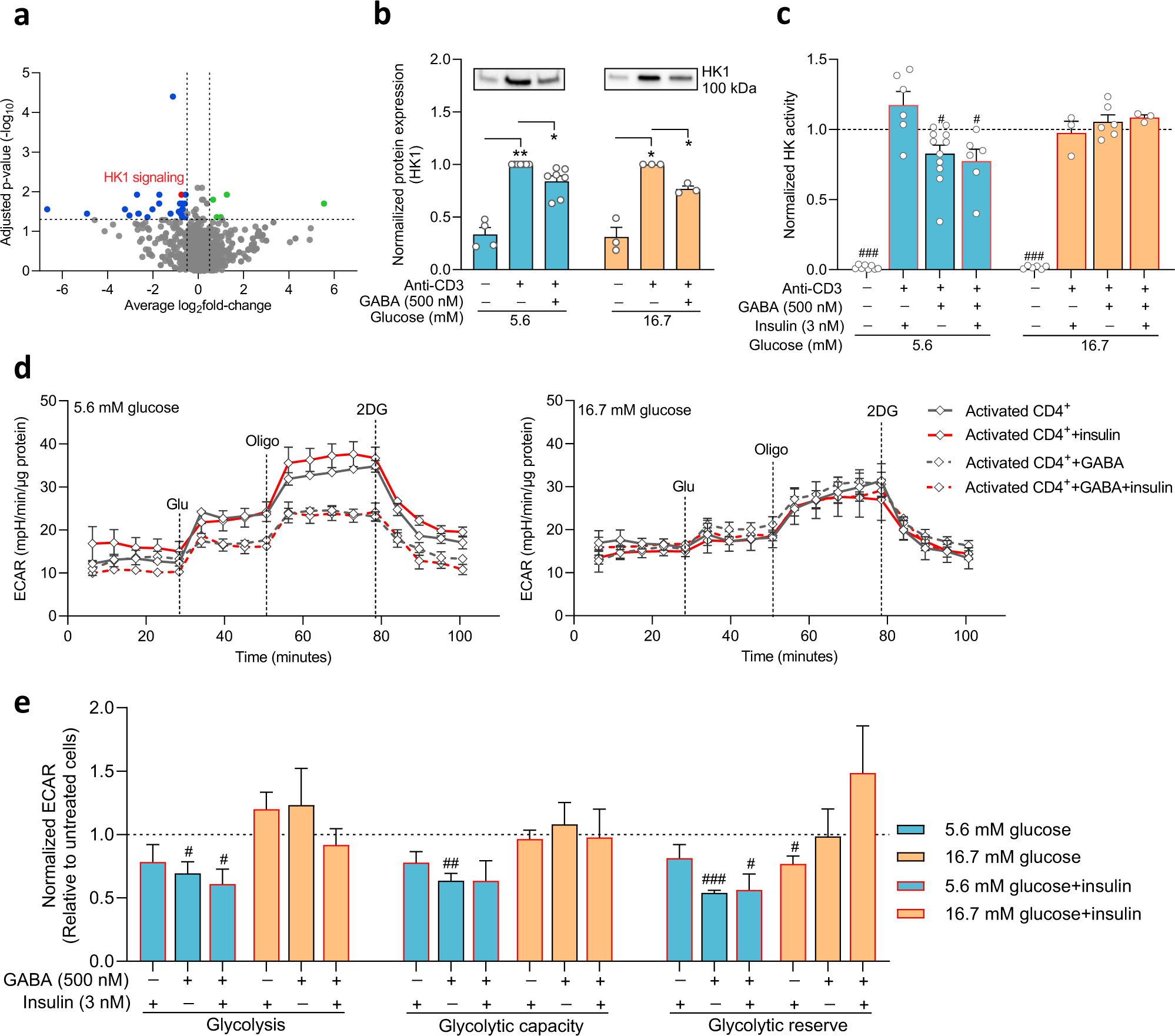
GABA inhibits hexokinase and glycolysis in activated CD4^+^ T cells. **a,** The volcano plot summarizes pathways that are significantly up-(green) or down-regulated (blue and red (HK1)) by GABA treatment in cells 72 h post activation (N=5), 5.6 mM glucose. The x-axis represents the average fold-change of all proteins measured by mass spectrometry within that pathway and every dot is one pathway. The y-axis represents the adjusted p-value (-log10 transformation). The horizontal line represents p = 0.05, and the vertical lines represent average log2 (fold change) < −0.5 or > 0.5. **b,** Western blot images and relative expression of HK1 protein (N=3-7). **c,** *In vitro* HK activity analyzed in resting and activated cells, 5.6 mM (N=12) or 16.7 mM (N=6) glucose. HK activity (µmol NADH/min/ml) was normalized to the activity of activated cells in the absence of drugs. **d,** Representative ECAR over time in cells 72 h post-activation treated with or without GABA (500 nM) or insulin at 5.6 mM (left panel) or 16.7 mM (right panel). Dashed lines indicate injections into media of glucose (Glu.), oligomycin (Oligo) or 2-deoxyglucose (2-DG). **e.** ECAR results (mpH/min/protein) were normalized to controls in absence of drugs for average glycolysis, glycolytic capacity and glycolytic reserve in cells 72 h post-activation treated with or without GABA or insulin, 5.6 mM (N=5) or 16.7 mM (N=5) glucose. Data are normalized to values of activated cells in the absence of drugs at each glucose concentration (**b, c, e**). Data represent mean ± SEM (**b, c, d, e**). Statistics: one sample t-test when compared to activated cell group (#, p < 0.05; ##, p < 0.01; ###, p < 0.001), repeated measures ANOVA or fitting a mixed-effects model followed by Tukey multiple comparison test. **N**: donors.

### Functional SGLT2 modulates glycolysis and cytokine release in activated CD4^+^ T cells

Facilitated glucose transport is the main route for glucose entry into activated T cells^3^. We confirmed the dominance of the facilitated transport on the cellular metabolic activity of activated CD4^+^ T cells (Fig. 6a) by examining the effects of antagonists of glucose transporters, WZB117, an inhibitor of the facilitated glucose transporters (GLUTs), and phlorizin, an inhibitor of Na^+^-dependent glucose transport (SGLT1 and SGLT2)^34^. WZB117 decreased whereas phlorizin had no effect at either glucose concentration on the metabolic activity of the T cells (Fig. 6a). In accordance, our RNAseq analysis showed several facilitated GLUTs gene transcripts (e.g., *SLC2A1* and *SLC2A3*) were prominently expressed in activated CD4^+^ T cells (Fig. 6b). Nevertheless, *SLC5A2*, the gene encoding the sodium-dependent glucose transporter SGLT2, was also detected at a modest level (Fig. 6b). Western blot and immunofluorescence staining data showed protein expression of SGLT2 in the cells (Fig. 6c, d).

**Figure 6.**
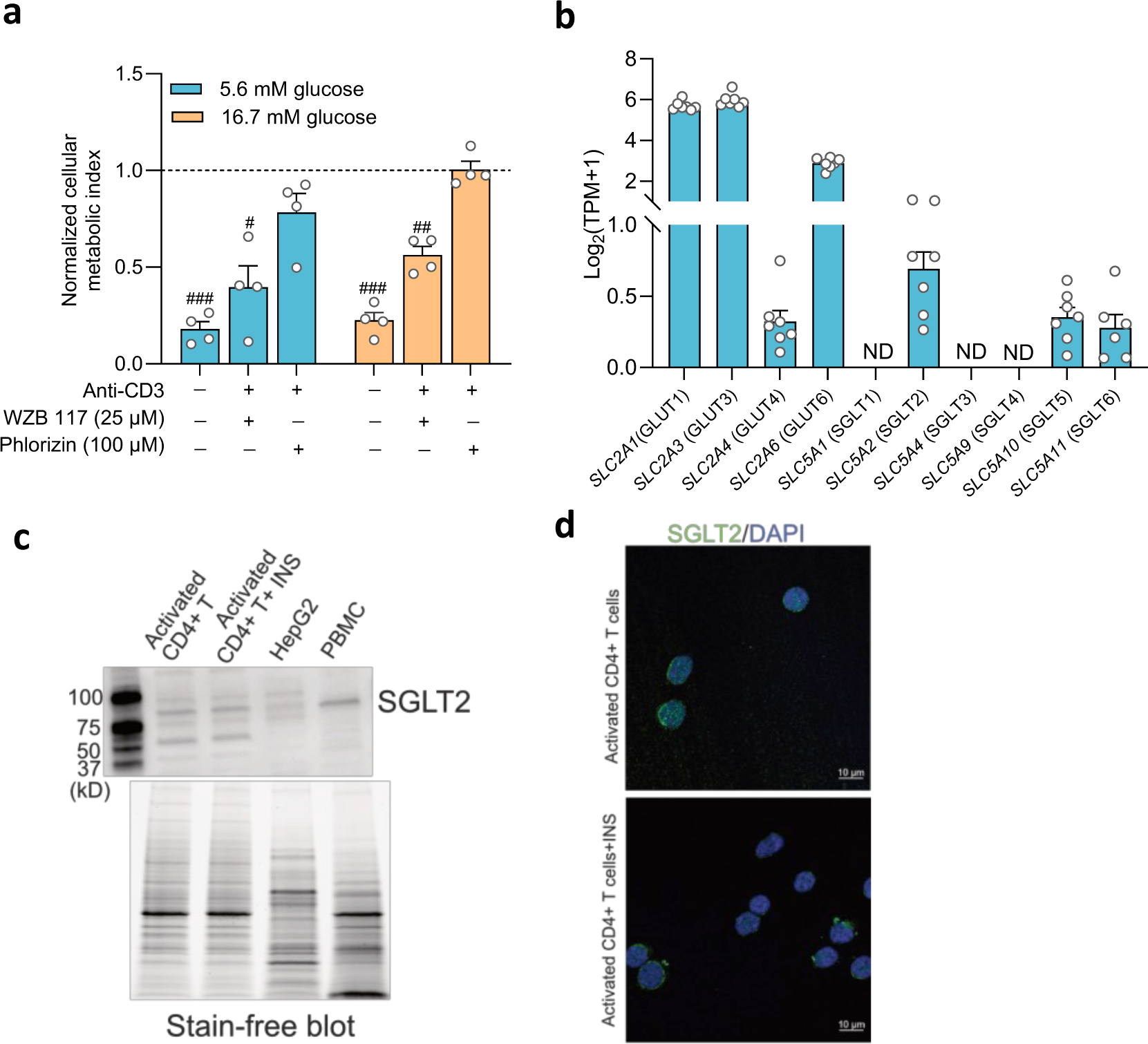
Sodium-glucose co-transporter 2 (SGLT2) is expressed in activated human CD4^+^ T cell. **a,** The cellular metabolic activity of resting and activated cells in presence of GLUT1 inhibitor, WZB 117 or SGLTs inhibitor, phlorizin was normalized to that of activated cells without drug treatment in 5.6 mM or 16.7 mM glucose (N=4). Data represent mean ± SEM. Statistics: one sample t-test when compared to activated cell group (#, p < 0.05; ##, p < 0.01; ###, p < 0.001) **b,** Relative gene expression levels of facilitated glucose transporters (GLUTs) and Na^+^-dependent glucose transporters (SGLTs) in activated cells at 5.6 mM glucose (N=6) determined by RNAseq analysis. TPM, transcripts per million. Data represent mean ± SEM. ND, not detected. **c,** Representative Western blot image of SGLT2 protein in activated CD4^+^ T cells and HEPG2 cell line. The stain-free blot image represents total proteins loaded for each sample, insulin (INS, 3 nM), 16.7 mM glucose. **d,** Representative immunofluorescence co-staining of SGLT2 (green) and DAPI (blue) in CD4^+^ T cells, 5.6 mM or 16.7 mM glucose. **N**: donors.

In presence of phlorizin or empagliflozin, another SGLT2 antagonist and a prescription medicine in type 2 diabetes, a significant reduction in glucose uptake was observed in insulin incubated cells (Fig. 7a). Phlorizin alone reduced glycolysis and glycolytic capacity at both 5.6 mM and 16.7 mM glucose (Fig. 7b), whereas insulin reduced or abolished the inhibition by phlorizin (Fig. 7b). Since phlorizin reduced glycolysis, it could be expected to reduce synthesis of biomolecules such as IFNγ and IL-10, similar effects to what we observed for GABA (see Figs. 1d, e, f). Indeed, phlorizin reduced release of IFNγ and IL-10 at both 5.6 mM and 16.7 mM glucose even in the presence of insulin (Fig. 8a, b) with the exception of IFNγ at 16.7 mM glucose with insulin, where only phlorizin together with GABA reduced IFNγ release, approximately 50%. In general, there was a trend for GABA to enhance the phlorizin reduction of release of IFNγ and IL-10. The results demonstrate functional Na^+^-dependent glucose carriers in CD4^+^ T cells and identify their relevance for production of cytokines.

**Figure 7.**
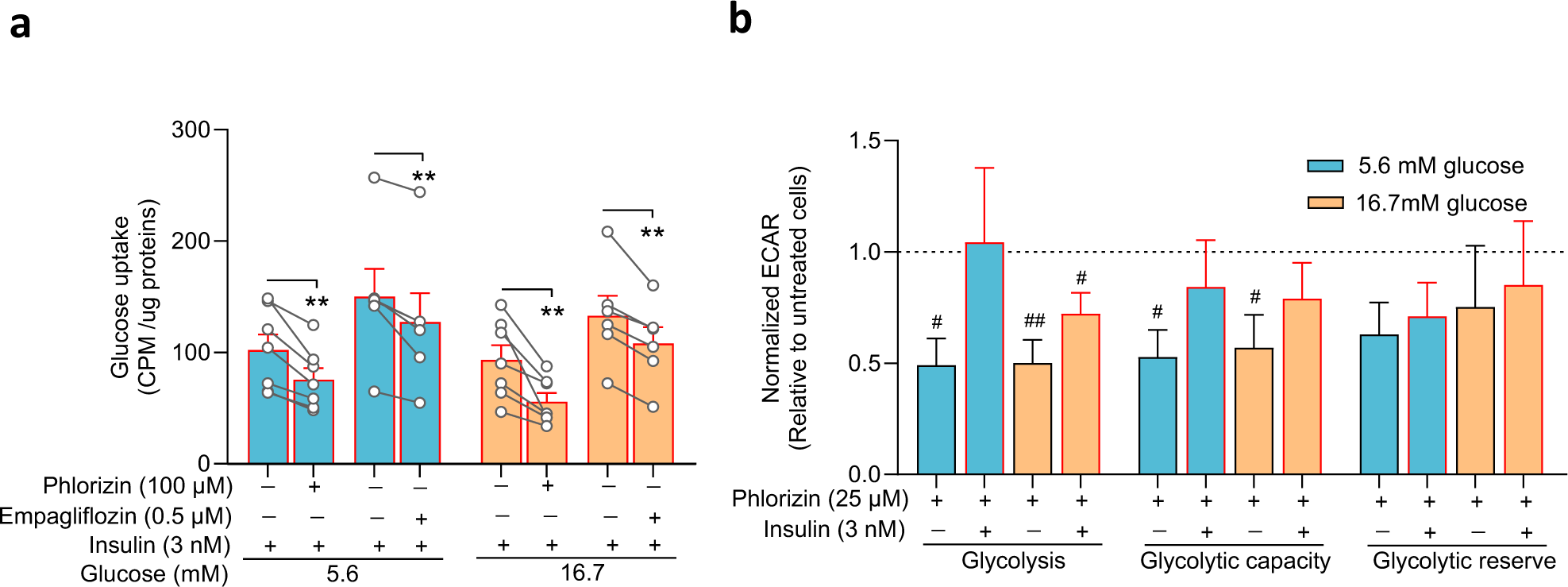
SGLT2 is functional and modulates glycolysis in activated human CD4^+^ T cell. **a,** Glucose uptake in absence or presence of phlorizin (N=7) and SGLT2 selective inhibitor, empagliflozin (N=6) in cells 72 h post-activation, with insulin, in 5.6 mM or 16.7 mM glucose. Data represent before and after inhibitor treatment from each donor as well as mean ± SEM. Statistics: two-tailed paired Student’s t-test, *, p < 0.05. CPM: count per minute. **b,** Extracellular acidification rate (ECAR) results (mpH/min/protein) normalized to activated cells in the absence of drugs for average glycolysis, glycolytic capacity and glycolytic reserve in cells, 72 h post-activation, incubated with phlorizin, with or without insulin in 5.6 mM (N=5) or 16.7 mM (N=5) glucose. Data represent mean ± SEM. Statistics: one sample t-test (#, p < 0.05; ##, p < 0.01). **N**: donors.

**Figure 8.**
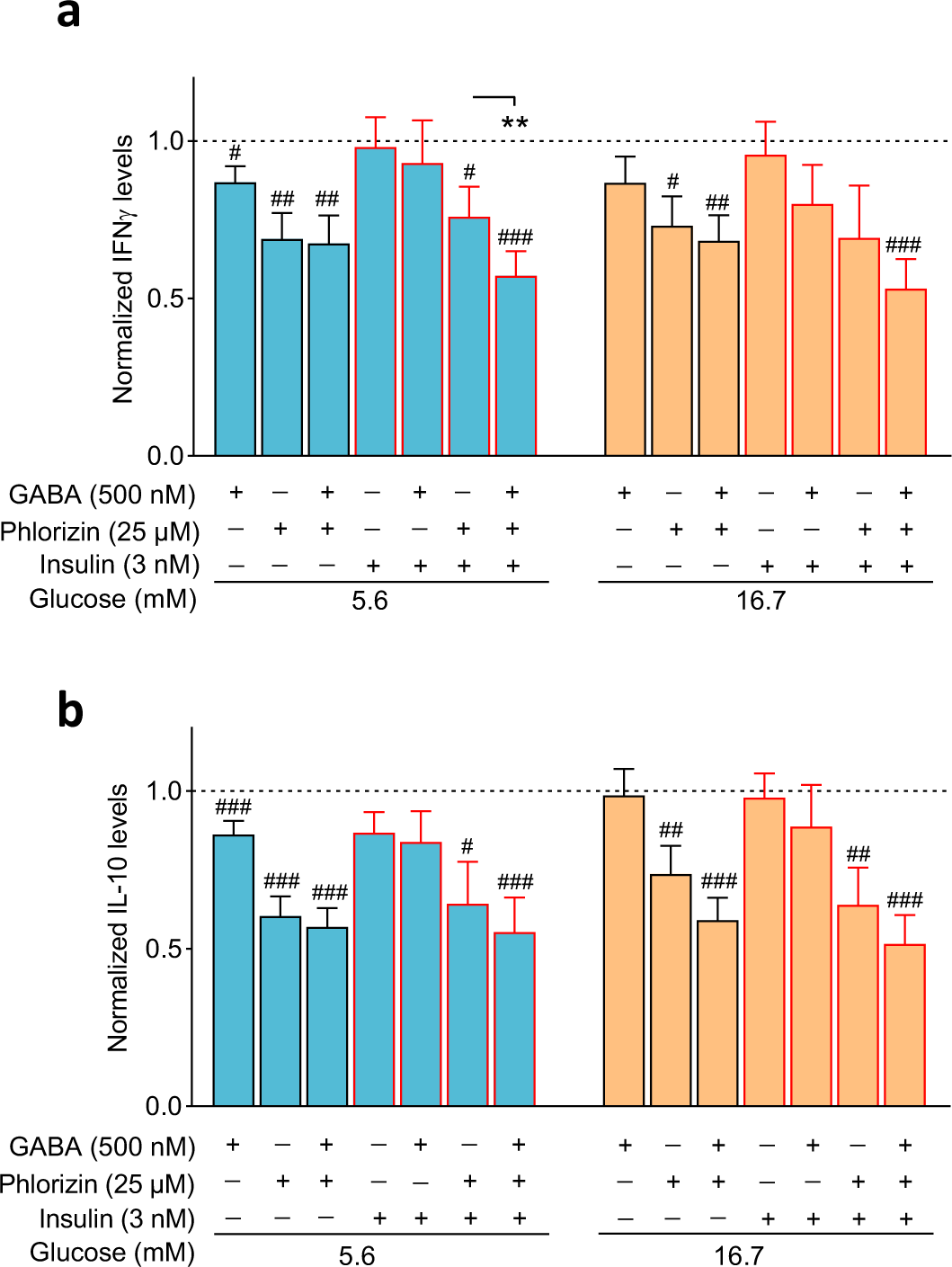
SGLT2 modulates cytokine release in activated human CD4^+^ T cell. Bar graphs show IFNγ **(a)** and IL-10 **(b)** release from activated cells 72 h post-activation in with or without GABA, phlorizin or insulin, at 5.6 mM or 16.7 mM glucose (N=17). Data represent mean ± SEM. Data are normalized to values of activated cells in the absence of drugs at each glucose concentration. Statistics: one sample t-test when compared to activated cell group (#, p < 0.05; ##, p < 0.01; ###, p < 0.001), repeated measure one-way ANOVA or Mixed-effects analysis with Tukey multiple comparisons, **, p<0.01. **N**: donors.

## Discussion

T cells modulate their metabolism to adapt to different environments^1,31,32^. When CD4^+^ T cells are activated, their metabolic activity is altered to meet the demands of cell growth, proliferation and effector functions. Here we revealed that GABA, glucose and insulin together orchestrate functions of CD4^+^ T cell, to-coordinate and to-set the activity level of distinct cellular pathways linked to glucose metabolism and that are central for T cells effector functions (Fig. 9).

**Figure 9.**
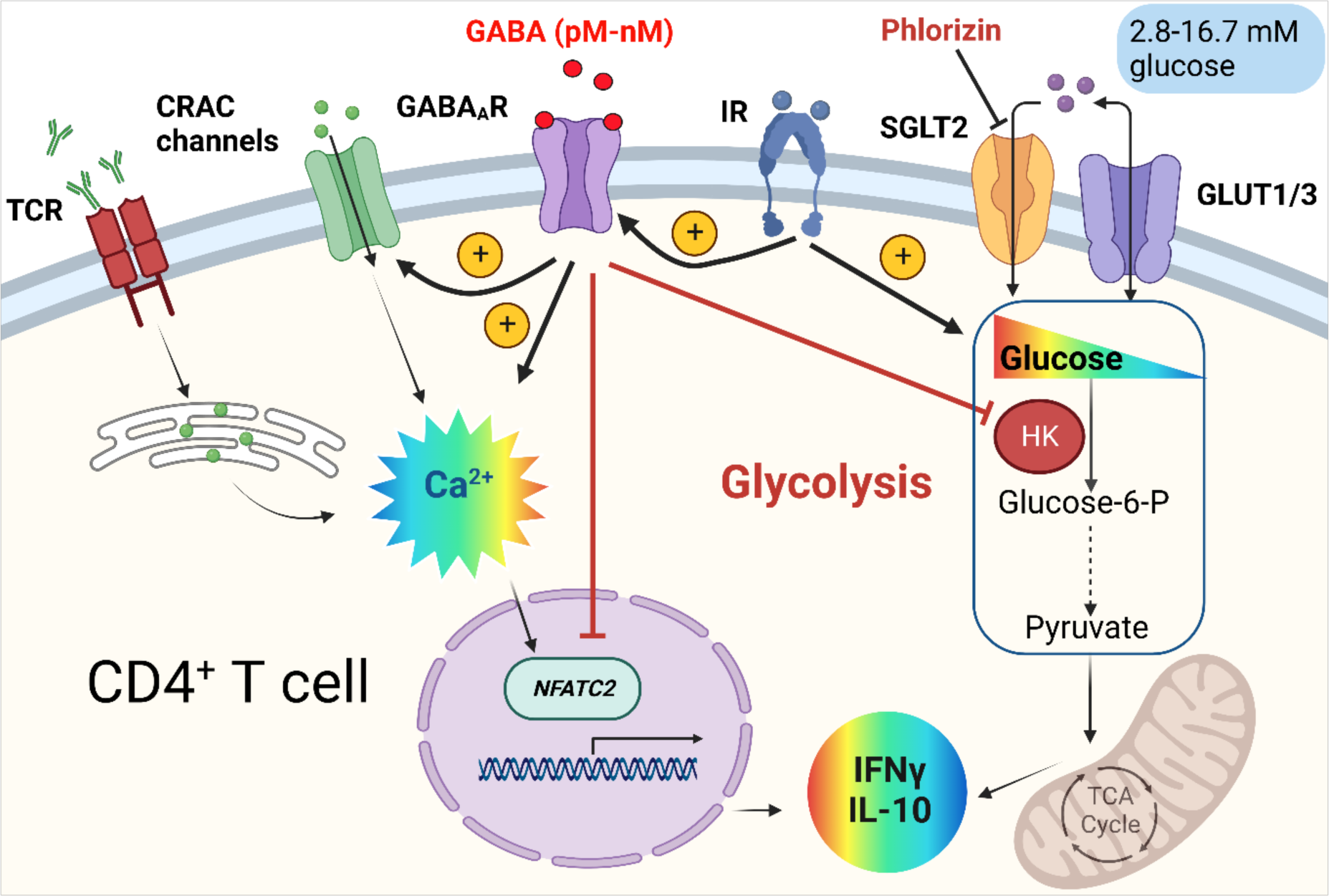
GABA, glucose and insulin regulate CD4+ T cell effector functions. In human CD4+ T cells, the activation of T cell receptor (TCR)/CD3 complex by anti-CD3 antibodies binding, initiates store-operated Ca2+ entry (SOCE) through Ca2+ release-activated Ca2+ (CRAC) channels. It regulates intracellular Ca2+ signals, drives changes in transcription factors, upregulates glycolysis and increases T cell effector functions. Physiological concentration of GABA (pM-nM) activates GABAA receptors (GABAAR) to modulate T cells metabolic activity and levels of cytokines (e.g., IFNγ and IL-10) by 1) enhancing function of CRAC channels and Ca2+ entry; 2) reducing expression of Nuclear Factor of Activated T cell 2 (NFATC2 or NFAT1); 3) decreasing expression and activity of hexokinase (HK), the 1st rate-limiting enzyme in aerobic glycolysis. In addition, insulin-insulin receptor (IR) signaling promotes glycolysis but paradoxically reduce glycolysis by enhancing GABAAR function. The fluctuation of blood glucose concentration affects glycolysis and cytokine production. Glucose uptake is mediated by not only the facilitated glucose transporters GLUT1/3 but also the sodium-dependent glucose transporter 2 (SGLT2). Phlorizin, a SGLT inhibitor, reduces glucose uptake, attenuates glycolysis and reduces IFNγ and IL-10 levels. Created with BioRender.com.

Proliferating T cells shift their metabolic phenotype to aerobic glycolysis at the expense of mitochondrial oxidative phosphorylation^31,35^. Although this switch is inefficient in terms of generating ATP, it confers advantage for the biosynthetic pathways that branch out from glycolysis^31,32^. GABA decreased cellular metabolic activity and release of proteins in a glucose concentration-dependent manner. Insulin shifted the GABA_A_R signaling by making it supersensitive to GABA and consequently enhanced Ca^2+^ entry through CRAC channels, an elegant way of modulating intracellular Ca^2+^ levels and intracellular processes affecting T cells effector functions.

CRAC channel-mediated SOCE regulates several Ca^2+^ signaling downstream pathways including the expression of facilitated glucose transporters (e.g., GLUT1) and glycolytic enzymes such as HK via activation of transcriptional factor NFAT in T cells^30^. Our data show that GABA decreases the expression of *NFATC2* mRNA transcript and the activity of HK2 at 5.6 mM glucose in activated CD4^+^ T cells. Consequently, at the level of HK there is a cross-over of GABA signaling and glucose transport for regulating glycolysis and, thereby, effector functions of T cells. Although glucose transport in CD4^+^ T cells is mainly mediated by facilitated glucose transporters, the present results identified significant Na^+^-dependent glucose (SGLT2) transport. This is highly relevant for the effector functions as only SGLTs can raise the intracellular glucose level above the extracellular ones. HK is the first enzyme in the glycolytic pathway and by converting glucose to glucose-6-phosphate (G-6-P), traps glucose within the cell^7^. Human HK has normally maximum activity in 5 mM glucose in the presence of natural inhibitors, such as G-6-P^36^, but HK activity is increased at glucose concentrations > 5 mM^36^. This is in line with our results, where GABA only succeeded decreasing HK enzymatic activity in 5.6 mM glucose. Na^+^-dependent glucose transport drives glucose into cells even at high intracellular glucose concentrations^34^. Glucose competes with G-6-P for the binding site on HK^36^, decreasing the inhibition of HK^36^, which results in increased substrate availability for biosynthesis as the intracellular glucose concentration increases^13^. Aerobic glycolysis directly regulates synthesis of IFNγ^37^ by engaging glyceraldehyde-3-phosphate dehydrogenase (GDPDH) and, thereby, relieving its inhibition of IFNγ mRNA translation. In concordance, activated CD4^+^ cells secreted more IFNγ in high glucose. GABA promoted a glycolytic switch to lower values for both glycolysis and the glycolytic capacity in 5.6 mM glucose that was gone in 16.7 mM glucose, whereas phlorizin was effective inhibitor at both glucose concentrations and, in particular, in the absence of insulin. Discovering that GABA_A_R activation together with SGLT2 inhibition can dramatically reduce the levels of IFNγ and IL-10, at both normal and high glucose concentrations, has implications for regulation of inflammation, as availability of biosynthetic intermediates-production fuels effector signaling molecules and functions^13^.

GABA_A_Rs are widespread in the brain and are present in many other tissues^38^ but GABA_A_Rs containing GABRR2 (ρ2 subunit) are rare outside the immune system, with the exception of the retina^39^. GABA_A_R-ρ2 might be a valuable and a rather specific drug target^28^. SGLT2 inhibitors such as empagliflozin, are already in clinical use^40^ and might be considered to cover new areas of application. Strengthening the GABA response and inhibiting SGLT2 in CD4^+^ T cells might be valuable tools^15,41,42^, when regulating inflammation is desirable.

In conclusion, physiological factors such as glucose, GABA and insulin guide the adaptation of CD4^+^ T cells and modify their effector functions. The reduction of metabolic activity and release of inflammatory molecules in response to GABA or SGLT2 inhibitors, revealed a switch that can be turned on or off. The study unveils the mechanisms of GABA modulation of CD4^+^ T cells and identifies a critical role for SGLTs in immune function. The results have implications for treatment of a number of diseases including diabetes but also maladies such as Covid-19 and cytokine storm, inflammatory diseases and cancer.

## Acknowledgements

We thank the following for support; The Swedish Children’s Diabetes Foundation, The Swedish Diabetes Foundation, The Swedish Research Council 2018-02952, EXODIAB, The Ernfors Foundation and The Thurings Foundation. We thank Dr Zhitao Gong for participation in some of the experiments and Universitetslektor Femke Heindryckx for the gift of the HepG2 cell line.

## Author Contributions

BB, ZJ, HH conceived project; HH, ZJ, AKB, SVK, BB, MKM designed experiments; HH, ZJ, AKB, SVK, OTR, SK, AIC and FA did experiments; ZJ, HH, AKB, SVK made figures; all authors analyzed results, BB wrote manuscript that was edited by ZJ, HH, AKB and then commented on by all authors.

## Supplementary information

**Supplementary Fig. 1:**
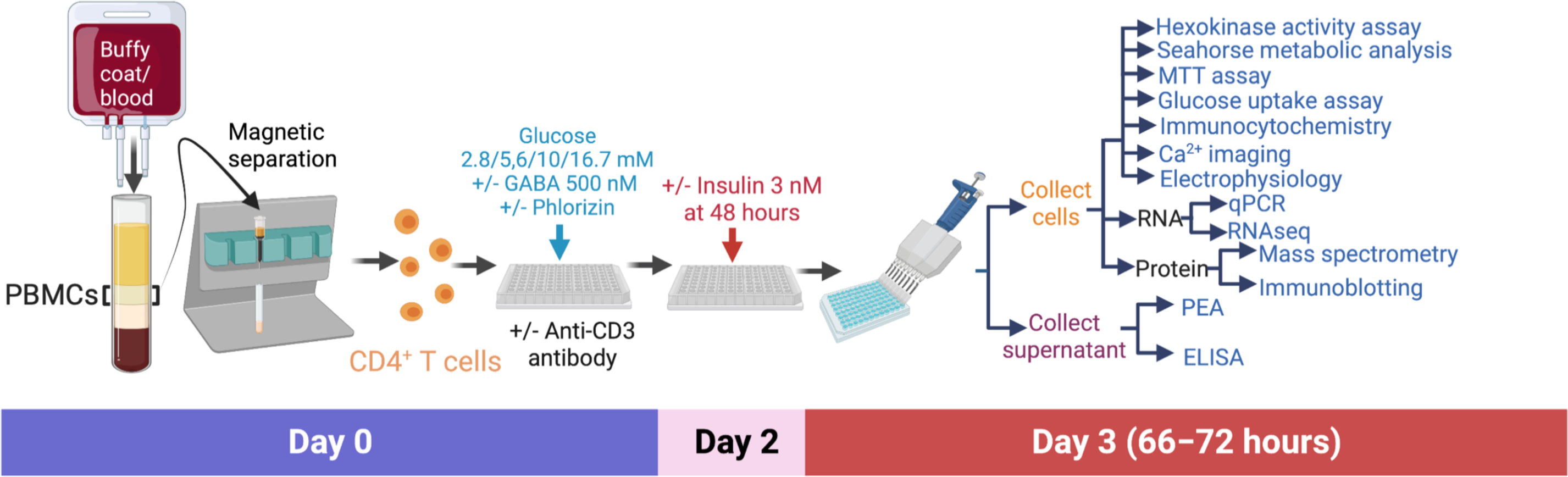
Schematic diagram of the experimental design. Created with BioRender.com.

**Supplementary Fig. 2:**
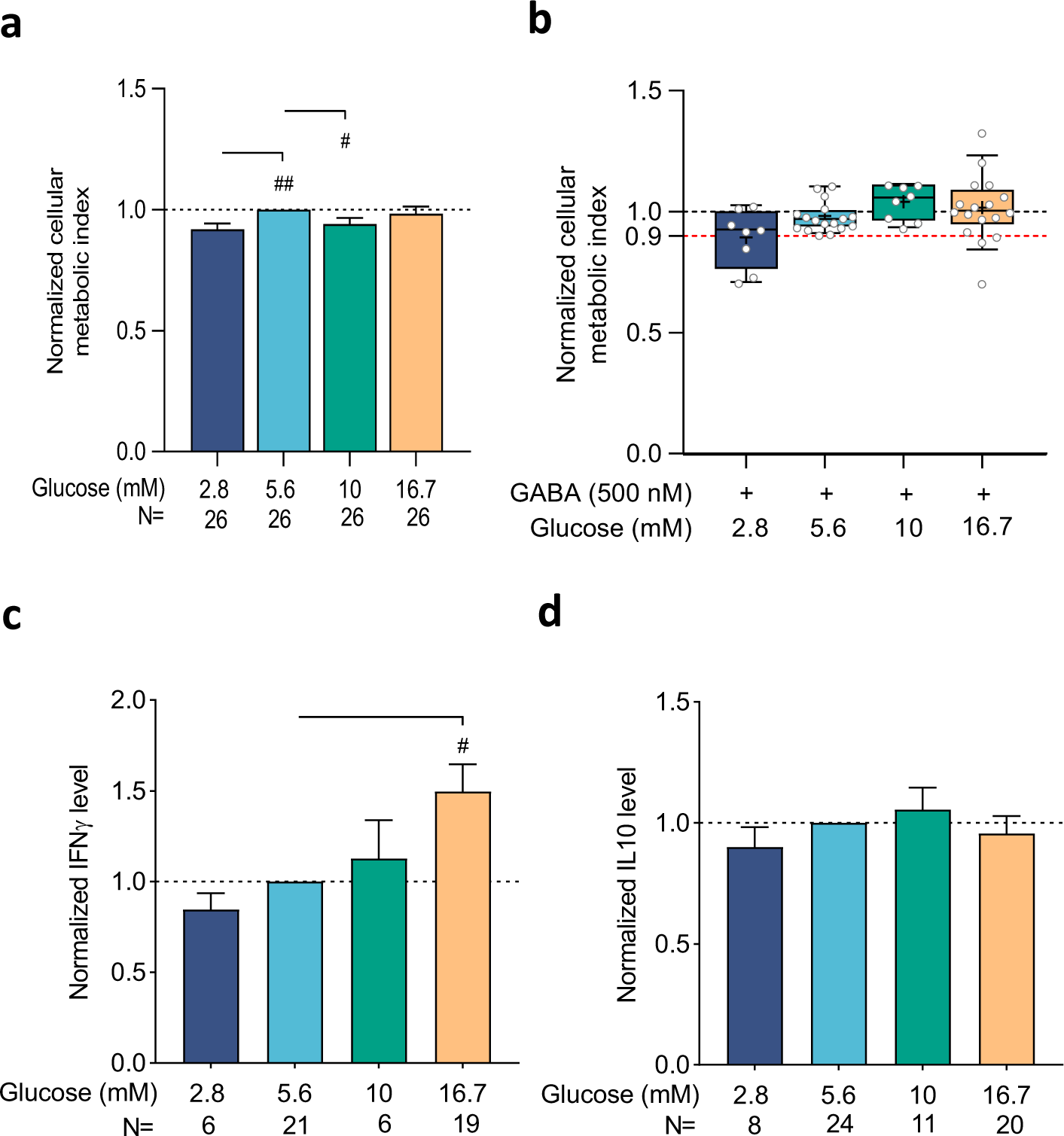
The effect of glucose on cellular metabolic activity and release of IFNγ and IL-10 from CD4^+^ T cells. **a,** Cellular metabolic activity of CD4^+^ T cells 72 h after activation in the presence of different glucose concentrations. **b,** Cellular metabolic activity from donors where <10% GABA inhibition at 5.6 mM glucose was observed. **c, d,** IFNγ (**c**) and IL-10 (**d**) released from CD4^+^ T cells 72 h after activation in presence of different glucose concentrations. Data are normalized to values at 5.6 mM glucose and presented as mean ± SEM. Statistics by one sample t-test when compared to 5.6 mM glucose group (#, p < 0.05; ##, p < 0.01), repeated measures ANOVA or fitting a mixed-effects model followed by Tukey test for multiple comparisons. N: donors.

**Supplementary Fig. 3:**
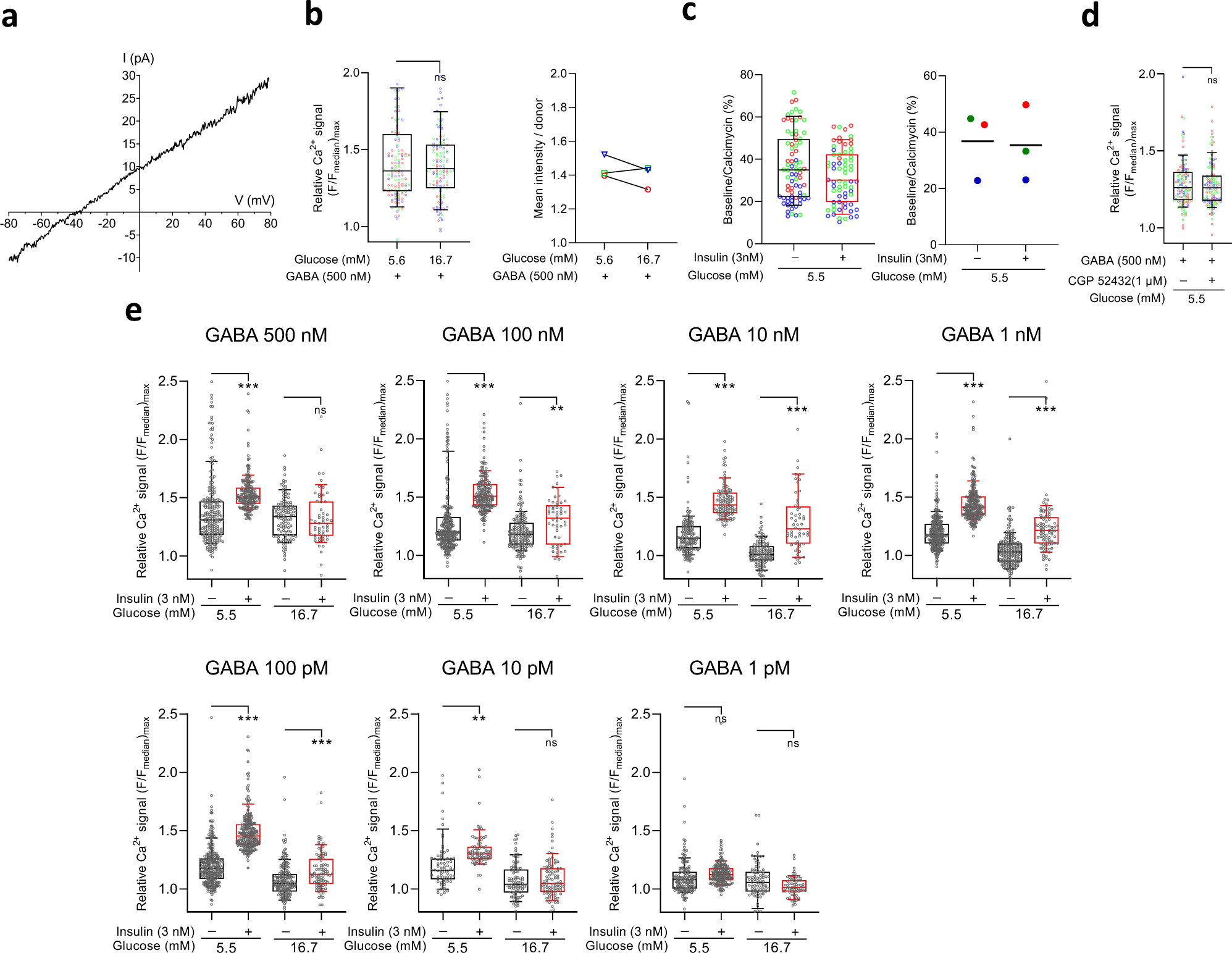
GABA-activated current and Ca^2+^ signaling in CD4^+^ T cells. **a**, A representative current-voltage relationship recorded in an activated CD4^+^ T cell in response to 500 nM GABA application by perforated patch-clamp, 5.6 mM glucose. **b,** GABA (500 nM)-activated relative Ca^2+^ signals (left panel) and mean intensity (right panel) recorded from activated CD4^+^ T cells (n=128 cells/ N=3 donors) in sequentially applied 5.6 mM and 16.7 mM glucose-containing media. Cells were activated in 5.5 mM glucose-containing media for 72 hours. **c,** The ratio of Ca^2+^ signals in the absence and presence of calcium-ionophore calcimycin (1 µM) in CD4^+^ T cells (n=85 cells left panel/ N=4 donors right panel) 72 h after activation with or without insulin treatment. **d,** GABA-activated relative Ca^2+^ signal recorded in CD4^+^ T cells (n=131 cells/ N=3 donors) in the absence or presence of GABA_B_ receptor antagonist CGP 52432 (1 µM) at 5.5 mM glucose. **e,** Relative Ca^2+^ signals induced by different concentrations of GABA in activated CD4^+^ T cells treated without or with insulin in 5.5 mM or 16.7 mM glucose. Statistics by two-tailed student’s t-test. ns, p > 0.05, **, p < 0.01; ***, p < 0.001.

**Supplementary Table 1:** Donors’ demographic information.

| Parameter | Normal Donors<br>(N=160) |
| --- | --- |
| Age (years) | 26.2 ± 0.25 |
| Sex (N, %) | 89 males (55.6%) |
|  | 71 females (44.4%) |

Data are presented as means ± SEM.

**Supplementary Table 2:**
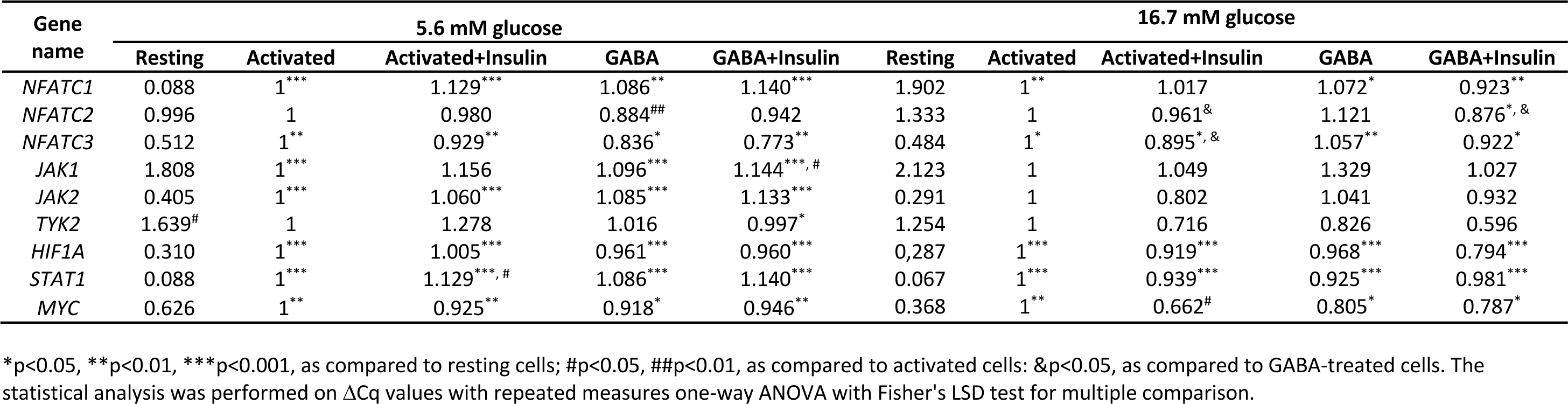
Relative gene quantification (normalized to activated cells) by qPCR (N=7-11 donors).

**Supplementary Table 3:** Reactome analysis of MS data (N=5 at 5.6 mM glucose). Proteins are named after UniProtKB.

**Top up-regulated proteins**
| Identifier | logFC | AveExpr | t | P.Value |
| --- | --- | --- | --- | --- |
| K2C79_HUMAN | 0.67266061 | 13.2926208 | 2.34394122 | 0.04308325 |
| CHM4A_HUMAN | 0.55827697 | 13.7149206 | 2.20925349 | 0.05380739 |
| HLAH_HUMAN | 1.03346983 | 14.8113954 | 2.03947592 | 0.07106636 |
| RAB2A_HUMAN | 1.37200668 | 14.1750267 | 2.03797883 | 0.07123998 |
| SC11A_HUMAN | 5.70132617 | 10.587487 | 1.91122932 | 0.08749562 |
| K22O_HUMAN | 0.48972535 | 12.2546633 | 1.88242372 | 0.09164997 |
| DYL1_HUMAN | 5.54957599 | 10.7818438 | 1.81112175 | 0.10274055 |
| PLXA4_HUMAN | 0.69449352 | 13.706242 | 1.7195177 | 0.11881622 |
| RL36_HUMAN | 6.02123532 | 12.32297 | 1.68555819 | 0.12534011 |
| IF4G2_HUMAN | 0.82748497 | 13.4787932 | 1.68221061 | 0.12600056 |

**Top down-regulated proteins**
| Identifier | logFC | AveExpr | t | P.Value |
| --- | --- | --- | --- | --- |
| TBB1_HUMAN | -7.55965074 | 8.58464831 | -2.74841088 | 0.02204275 |
| STA5B_HUMAN | -7.60692039 | 6.04448902 | -2.44305026 | 0.03656351 |
| EXOS3_HUMAN | -6.6014009 | 5.67159622 | -2.16624065 | 0.05775108 |
| BGAL_HUMAN | -6.67048214 | 5.77846898 | -2.13711593 | 0.06057899 |
| AL1B1_HUMAN | -6.67577438 | 5.87276979 | -2.08793829 | 0.06566002 |
| HXK1_HUMAN | -0.73989819 | 14.3697968 | -2.06803575 | 0.06783079 |
| HLAG_HUMAN | -6.34860408 | 5.64345434 | -2.03980339 | 0.07102844 |
| COPG1_HUMAN | -5.15586276 | 10.0461233 | -1.80136877 | 0.10435116 |
| BCL2_HUMAN | -4.91388319 | 9.86964223 | -1.73134501 | 0.11661798 |
| PIP_HUMAN | -4.68190741 | 9.67505331 | -1.66669489 | 0.12910318 |

## Methods

### Study individuals and collection of samples

Human blood buffy coat and blood samples were obtained from Uppsala University Hospital (Uppsala Akademiska sjukhuset), protocol approved by the Regional Research Ethical Committee in Uppsala. All donors were voluntary recruited and informed consent were signed. In total, samples from 160 normal donors, 89 men and 71 women with mean age 26.2 ± 0.25 years were included in the study.

### Chemicals and reagents

Drugs, buffers and salts unless otherwise specified were purchased from Sigma-Aldrich/Merck (Steinheim, Germany). TACA, muscimol, TPMPA, YM-58483, WZB 117 were purchased from Tocris (UK, 7A/109031). Phlorizin and empagliflozin were obtained from AdooQ Bioscience. Urea and NH_4_HCO_3_ were from Acros Organics, Geel, Belgium and trypsin and dimethyl sulfoxide (DMSO) from Thermo Scientific (Waltham, MA). Tris(hydroxymethyl) aminomethane hydrochloride (Tris), LC-MS grade water, acetonitrile and formic acid were obtained from Fisher Scientific (Geel, Belgium).

### CD4^+^ T cells isolation and activation

Peripheral blood mononuclear cells (PBMCs) were isolated by density-gradient centrifugation using Ficoll-Paque™ PLUS (Sigma Aldrich, Sweden). CD4^+^ T cells were then purified from the isolated PBMCs by negative selection using humanCD4^+^ T cell isolation kits (Miltenyi Biotec, Germany) according to the manufacturer’s instructions. The freshly isolated CD4^+^ T cells (1x 10^6^/ml) were suspended in RPMI 1640 glucose-free medium (Gibco, Fisher Scientific, Sweden) supplemented with 2 mM glutamine, 10% heat-inactivated dialyzed fetal bovine serum, 100 U/ml penicillin, 10 mg/ml streptomycin, and 5 μM β-mercaptoethanol (Gibco, Fisher Scientific, Sweden). The activation of the cells was performed in either 96-well (10^5^ cell/well) or 24-well (10^6^ cell/well) plates based on the purpose of the experiment. CD4^+^ T cells were activated with 3 μg/ml plate-bound anti-CD3 (BD Biosciences, USA, 555329) in the presence of different concentrations of glucose (2.8 mM, 5.6 mM, 10 mM and 16.7 mM) for 66^_^72 h and indicated drugs were added based on the experimental design.

### MTT Assay

The metabolic activity of CD4^+^ T cells was assessed in 96-well plates with colorimetric MTT (3-(4,5-Dimethylthiazol-2-yl)-2,5-Diphenyltetrazolium Bromide) assay. It is commonly used to measure cellular metabolic activity based on mitochondrial dehydrogenase in living cells as an indicator of cell viability and proliferation^1^. After 66^_^72 h of activation, a PBS-soluble tetrazolium dye MTT (final concentration 1mM) was added and incubated for additional 4 h at 37 °C. Thereafter, the cells were centrifuged at 2000 rpm for 10 min to concentrate the insoluble purple formazan pellets. The supernatants were collected and stored at −80 °C for later biomolecules analysis. The formazan crystal pellets were dissolved in DMSO and the plate was read 10 min later using Multiskan FC Microplate Photometer (Thermo Fisher, USA) at 550 nm.

### Multiplex Proximity Extension Assay (PEA)

The supernatants collected at the end of the MTT assay were sent on-ice to the SciLifeLab Affinity Proteomics Unit in Uppsala, Sweden, for analysis. Inflammatory proteins were measured using high-throughput multiplex PEA with an Olink Target 96 Inflammatory panel (OlinkProteomics, Uppsala, Sweden). PEA measures simultaneously 92 proteins/biomolecules in different sample matrices where it combines a dual-detection immunoassay and quantitative PCR. Normalized protein expression (NPX) values, an arbitrary unit on the log2 scale, were used to present the data. A protein expression (NPX value) is considered acceptable if the value is above the limit of detection (LOD) for each particular protein. Proteins that were detected in more than 75% of the samples were analyzed in this study: 59 out of 92 proteins had detectable levels. Data for resting, activated and GABA-treated activated CD4^+^ cells supernatants were analyzed separately. The biomolecules in the supernatants were assessed for each donor at each glucose concentration. The data was computed, processed and analyzed with support from NBIS (National Bioinformatics Infrastructure Sweden). The log2FC (FC =fold change) is the average increase in NPX in GABA as compared to activated alone, this value is both estimated based on the linear mixed effects model and computed as the mean over the GABA-activated difference and plotted in heatmaps. Paired t-test was used followed by multiple testing correction using false discovery rate (FDR) method (Two-stage step-up method of Benjamini, Krieger and Yekutieli) with a threshold 12%.

### Determination of IFNγ and IL-10 concentrations in supernatants

The concentrations of IFNγ and IL-10 were measured with enzyme-linked immunosorbent assay (ELISA). The supernatants were diluted when necessary to be in the optimal linear optical range and analyzed using commercial kits for IFNγ (Human IFN-gamma DuoSet ELISA, R&D Systems, USA and Bio legend, UK), and IL-10 (Biolegend, UK) according to manufacturerś instructions. Optical densities were measured using FC Microplate Photometer (Thermo Fisher, USA) at 450 nm, and 540 nm was used as a correction wavelength. The levels of IFNγ or IL-10 were normalized to controls (activated CD4^+^ T cells in the absence of drugs).

### Total RNA and protein extraction and, bulk RNA sequencing

Total RNA and protein were extracted from activated CD4^+^ T cells, in the absence or presence of 500 nM GABA, using RNA/Protein Purification Plus Kit (Norgen Biotek, Ontario, Canada). During the extraction process, total RNA was treated with RNase-Free DNase I Kit (Norgen Biotek, Ontario, Canada) to eliminate contamination of genomic DNA. The extracted proteins were quantified using DC protein assay kit (Bio-Rad, USA), and immediately frozen at −80°C. Total RNA samples were submitted to the GENEWIZ NGS facility (Germany) and quality control analysis showed that all RNA samples had an RNA integrity number (RIN) >9 (9.48 ± 0.1, mean ± SEM). Strand-specific RNAseq cDNA libraries with polyA selection were generated and sequenced using Illumina HiSeq sequencing platform (12 samples, 20.8-31.1 million reads, 150 bp paired end per sample). Sequence reads were filtered and trimmed using Trimmomatic V.0.36 to remove low quality reads. The trimmed reads were further mapped and aligned to the Homo Sapiens GRCh38 reference genome (ENSEMBL) using the STAR aligner v.2.5.2b. Unique gene hit counts were calculated using feature Counts from the Subread package v.1.5.2.

### Real-Time Quantitative Reverse Transcription (qPCR)

The cDNA was synthesized using SuperScript reverse transcriptase IV (Invitrogen Thermo Scientific, USA) in a 20 μl reaction following the standard protocol from manufacturer. The qPCR reaction was then performed using qPCRBIO SYBR or Probe Mix low-ROX (PCR Biosystems, UK; PB20.11-05), and gene-specific primers (Table S1) or Taqman probes (*GABRR2* Hs00266703_m1; *IPO8* Hs00914057_m1, ThermoFisher Scientific, USA). Changes in the abundance of each transcript were normalized to the expression of reference genes (*IPO8*). Relative expression levels were calculated by normalizing to non-treated activated CD4^+^ T cells obtained from the same donor at the same glucose concentration using 2^-ΔΔCt^ method. All samples were run in duplicate. The real-time qPCR amplification was performed in 10 μl reaction mix in a 384-well plate using a QuantStudio™ 5 instrument (ThermoFisher Scientific). All runs started with an initial denaturation step of 2 min at 95 °C, followed by 45 cycles of 95 °C for 5 s and 60 °C for 30 s.

### Mass spectrometry (MS)-based bottom-up proteomics

The lysates were purified and tryptic digested for MS-based bottom-up proteomic analysis using Filter Aided Sample Preparation (FASP)^2, 3^. Briefly, 20 μg of total protein were transferred onto centrifugal filter units (Microcon-30 kDa; Merck, Darmstadt, Germany) and washed with a buffer containing 8 M urea and 100 mM Tris (pH 8.5). The washing step was repeated after reduction with 8 mM dithiothreitol (DTT), alkylation with 50 mM iodoacetamide (IAA) and removal excess of IAA with 8 mM DTT. Before tryptic digestion (enzyme-to-protein ratio 1:50 (w/w)) the filter was washed three times with NH_4_HCO_3_. After incubation in a wet chamber at 37 °C for 16 h the resulting peptides were washed twice from the filter by adding 50 mM NH_4_HCO_3_. Trifluoroacetic acid was then added to a final concentration of 1% (v/v) and the samples were dried at 45 °C. Finally, the samples were reconstituted in 3% acetonitrile and 0.1% formic acid in water to a final concentration of 150 ng protein/μl. The tryptic peptides were analyzed on a nanoAcquity UPLC system equipped with a C18, 5 μm, 180 μm x 20 mm trap column and an HSS-T3 C18 1.8 μm, 75 μm x 250 mm analytical column (Waters Corporation, Manchester, UK) coupled to a Synapt G2 Si HDMS mass spectrometer with an electrospray ionization source (Waters Corporation, Manchester, UK). The UDSM^E^ approach with positive ionization was used^2, 3^. Mobile phase A contained 3 % DMSO and 0.1 % formic acid in water and mobile phase B 3 % DMSO and 0.1 % formic acid in acetonitrile. After loading 300 ng of protein on the column in trapping mode, peptide separation was performed with a gradient run from 3 - 40% (v/v) mobile phase B over 120 min. The flow rate was set to 0.3 μl/min and the column oven was set to 40 °C. Method performance was controlled with a commercially available HeLa digest (Thermo Scientific, Waltham, MA). UPLC-MS data processing and label-free quantification: ProteinLynx Global Server (version 3.0.3, Waters Corporation, Milford, MA) was used for raw data processing. With FDR of 10%, a database search against a randomized UniProt human database (Uni-ProtKB version 14/01/2020) was performed. Carbamidomethyl cysteine was set as fixed modification, acetyl lysine, C-terminal amidation, asparagine deamidation, glutamine deamidation and methionine oxidation as variable modifications and trypsin as digest reagent. Minimum peptide matches per protein were 2 and minimum fragment ion matches per peptide and protein were 1 and 3, respectively. For label-free quantification analysis, the identified proteins were further processed using ISOQuant 1.8 as described elsewhere^2^. Following the TOP3 quantification approach the average intensity of the three most intense peptides of each protein were used for relative protein quantification. ISOQuant settings are provided in Table S2.

### Immunoblotting

CD4^+^ T cells were lysed in RIPA buffer (Sigma-Aldrich, R0278) containing protease inhibitor cocktail (Roche, 11 836 153 001) and phosphatase inhibitors (Sodium orthovanadate, Sigma-Aldrich, S6508). Proteins were heated at 100 °C for 5 min in SDS loading buffer (275 mM Tris. HCL, 9% SDS, 50% glycerol, 0.03% bromophenol blue) containing 9% beta-mercaptoethanol and separated by Mini-Protean TGX gel 4 - 15% (BioRad, stain-free gels #4568084, 4561035) and blotted onto Amersham Hybond-P PVDF membrane. Membranes were blocked with blocking buffer (EveryBlot BioRad, 12010020) for 20 min at room temperature (RT) and incubated overnight at 4 °C with the following primary antibodies: HK1 (GenTex, GTX105248, 1:3000), SGLT2 (ProteinTech, 24654-1-AP, 1:500) and GABRR2 (Invitrogen, PA5-41008, 1:1000). After immunoblotting, membranes were washed in TBST and incubated with Peroxidase-conjugated AffiniPure secondary antibodies (Jackson ImmunoResearch, 111-035-045, 1:5000). Protein detection was done using chemiluminescent ECL reagent (Thermo scientific, 32209). Quantitative densiometric analysis of the immunoblotted bands was performed using Image Labs software (BioRad Laboratories, V6.0.0) and data were normalized for each sample using stain-free technique.

### Immunofluorescence staining

CD4^+^ T cells were harvested after 72 h of anti-CD3 activation on cytoslides (Thermo scientific, 5991056) using a Shandon cytospin centrifuge (400 rpm for 4 min at RT). After drying, cells were fixed with 4% PFA in PBS for 15 min and washed 3 times, followed by membrane permeabilization with 0.1% Triton-X for 5 min. Un-specific binding was blocked with 5% bovine serum albumin and 10% donkey serum (Jackson ImmunoResearch, 017-000121) in 1X PBS for 60 min. Cells were then washed and incubated with primary antibodies diluted in blocking solution overnight at 4 °C followed by incubation with an AlexaFluor 488-conjugated AffiniPure Donkey Anti-Rabbit IgG secondary antibody (Jackson ImmunoResearch, 711-545-152, 1:500) for 1h at RT. Finally, cells were washed again and counterstained with DAPI for 5 min and slides were mounted in ProLongTM Gold antifade reagent (Invitrogen). Images were acquired using confocal microscopy (LSM700, Zeiss) with 63X objective. For statistics, ≥ 50 cells from 3 donors were quantified using ImageJ (NIH) software. Primary antibodies and concentrations included rabbit anti-insulin receptor, InsR (Invitrogen, PA5-29286, 1:800), rabbit anti-GABRR2 (Invitrogen, PA5-41008, 1:600), and rabbit anti-SGLT2 (ProteinTech, 24654-1-AP, 1:100).

### *In vitro* Hexokinase activity assay

Hexokinase activity was measured at RT indirectly as a readout of NADH production, a by-product of the reaction converting glucose-6-phosphate to 6 phosphogluconolacton by the enzyme glucose-6-phosphate dehydrogenase (GPDH). The cells were harvested, centrifugated and washed twice with PBS. Washed cells were resuspended in 1 ml assay buffer solution, on ice, included in the hexokinase enzymatic assay kit (Bio Vision, K789-100). After centrifugation at 4 °C for 5 min, the resulting supernatant was used as the final cell extract for the enzymatic assay.

### Electrophysiological recordings

The electrophysiological recordings from the isolated CD4^+^ T cells were done using the perforated whole-cell patch-clamp configuration. In order to keep the intracellular chloride concentration intact, gramicidin was used to obtain the perforated configuration. Gramicidin was diluted in DMSO at the stock concentration 2 mg/ml and the final concentration of the gramicidin in the pipette solution was 2.6 μg/ml. The pipette solution with the gramicidin was protected from light, kept cold and used within 2 hours after the preparation. After making the cell-attached configuration, the access resistance was monitored until characteristic slow and low-amplitude capacitance transients were observed and the access resistance decreased from GΩ range to the range of tens of MΩ. The extracellular solution (in mM) contained: 157 NaCl, 4.5 KCl, 0.5 CaCl_2_, 1 MgCl_2_, 5 HEPES and 5.6 or 16.7 glucose (pH 7.4, adjusted with NaOH). The pipette solution consisted of (mM): 149 KCl, 2 CaCl_2_, 1 MgCl_2_, 1 NaCl, 10 HEPES, 5.6 or 16.7 glucose (pH 7.3 adjusted with KOH). The patch-clamp pipettes were made from borosilicate glass and had a resistance of 12–14 MΩ when filled with the pipette solution and immersed in the extracellular solution. Recordings were done using Axopatch 200B amplifier, filtered at 2 kHz and digitized on-line at 10 kHz using an analog-to-digital converter. To record the electrophysiological data Clampex 10.5 (Molecular Devices, San Jose, CA, USA) software was used. Currents were recorded at +30 mV or by a ramp protocol, where the potential was changed from −80 to +80 mV in 1 s, the holding potential (Vh) was kept −40 mV. The single sweep duration was 20 s with a break between sweeps 5 s. After application of a drug, the chamber was perfused 2-6 min with extracellular solution only before next application of a drug. Currents in the absence of drugs or in the presence of picrotoxin were subtracted from the test-drug current response.

### Live cell time-lapse real-time Ca^2+^ imaging

Cells were loaded with a calcium indicator, 3 μM Fluo-8H AM (AAT Bioquest, Pleasanton, CA) for 15 min at 37 °C and seeded on 5% 3-aminopropyltriethoxysilane-coated coverslip for 15 min at 37 °C. Time lapse images were acquired by confocal microscopy (LSM700, Zeiss) with 40X objective at an interval of 1-1.5 second per image. Drugs were diluted in RPMI 1640 without phenol red and perfused at indicated time and concentration by a peristaltic pump. After application of each drug, cells were perfused with the extracellular solution (RPMI 1640) before next application. Area for single cells was marked and absolute fluorescence intensity values (F) were extracted using ZENBlue software. Relative intensity (F/Fmedian) for each cell was calculated for every recorded image and plotted against time. Further, maximum relative intensity of the cells under drug application time is considered as drug’s response. Data is represented for all cells and as box-whiskers plot.

### Metabolism Assay

Isolated CD4^+^ T cells were plated in a XF^e^96 cell plates that were coated with Cell-Tak (#354240, Corning, New York, USA) according to manufacturer’s protocol. The metabolic function of activated CD4^+^ T cells cultured for 3 days *in vitro* was analyzed by measuring the extracellular acidification rate (ECAR) using an XF^e^96 extracellular flux analyzer (Seahorse Bioscience). The cells were kept in XF media (Seahorse Bioscience) supplemented with 5 mM glucose (Sigma Aldrich), and subjected to glycolysis stress-test protocol using sequential injection of 25 mM glucose, 2 μM oligomycin, and finally 15 mM 2-deoxyglucose (2DG). Measurements for each experiment were conducted at least in triplicates and glycolysis, glycolytic capacity, and glycolytic reserve calculated and values normalized to protein content ^4, 5^.

### Glucose uptake assay

2x 10^6^ activated CD4^+^ T cells (72 h activation) with insulin treatment from 48 h after activation were pelleted, resuspended in glucose-free RPMI-1640 media, incubated for 10 min in the incubator at 37 °C, and washed once with the same media. Glucose-free media (500 μl) containing 1μCi/ml [^14^C] glucose (Perkin Elmer) were added to cells in the absence or presence of phlorizin (100 μM) or empagliflozin (0.5 μM). Cells were incubated for 40 min at 37 °C, pelleted by centrifugation at 300x g for 5 min and washed three times in cold PBS. Cells were then lysed in RIPA buffer and ^14^C content was measured with Gold XR scintillation fluid using a liquid scintillation counter (Perkin Elmer). The glucose uptake was calculated by normalizing counts to protein contents for each sample.

### Statistical analysis

Statistical analysis was performed using GraphPad Prism 9 GraphPad Prism (GraphPad Software Inc., La Jolla, CA, Version 9.3.1). Normalized PEA data were assessed by two-tailed paired t-test followed by Benjamini-Hochberg’s FDR (False Discovery Rate) method. The significance level is set to 10% FDR. Cellular metabolic activity, cytokine levels, mRNA, protein expression and ECAR of CD4^+^ T cells in the presence of indicated drugs were normalized to controls (activated cells without drug treatment) from each donor and for each glucose concentration. Normalized data were analyzed by one-sample t-test when compared to controls. A comparison between paired groups was performed using repeated measures or mixed-model effects one-way ANOVA followed by Tukey or Fisher’s LSD multiple comparisons tests. Unpaired multiple-group comparison was performed using ordinary one-way ANOVA followed by Tukey test. Changes in protein levels in MS data were analyzed using Quantitative, multi-dataset Pathway Analysis (ReactomeGSA, https://reactome.org/PathwayBrowser/#TOOL=AT)^6^ and volcano plot was subsequently generated. Error bars in the bar graphs represent SEM unless otherwise specified and the statistical significance level was set to 0.05.

